# Proteome analysis of daily urine samples of pregnant rats unveils developmental processes of fetus as well as physiological changes of mother rats

**DOI:** 10.1101/2025.02.14.638372

**Authors:** Haitong Wang, Linna Ge, Sijie Chen, Longqin Sun, Wei Sun, Youhe Gao

## Abstract

Significant physiological changes occur in both fetus and mother during pregnancy. Urine proteins have been shown to reflect a wide range of physiological and pathological changes in the body. To comprehensively explore the daily changes in urine proteins during pregnancy, this study employed low-abundance protein-enriched magnetic nanobeads to conduct an in-depth analysis of the daily changes in urine proteins throughout the entire pregnancy of rats. Based on the 3,455 identified urine proteins, fetal and maternal dynamic changes were observed in the pregnancy group compared to the control group, including blastocyst formation and cell division in the early stage of pregnancy, embryonic development and organ morphogenesis in the intermediate and mid-to-late stages of pregnancy, and maternal-specific change such as lactation in the late stage of pregnancy. These results indicate that urine protein can reflect the fetal and maternal dynamic physiological alterations during pregnancy, which suggests the potential value of urine protein analysis in pregnancy health monitoring. It is emphasized that the analysis focuses on the daily variations in the urine proteins, as these daily changes are expected to reveal more dynamic and detailed information about the physiological processes during pregnancy.

## 1. Introduction

Pregnancy is an extremely complex and precisely regulated physiological process, during which a series of significant changes occur in both the mother and the fetus. The maternal physiological systems need to make adaptive adjustments to meet the demands of fetal growth and development, while the fetus undergoes a gradual developmental process from a fertilized egg to a complete individual in the uterus [1, 2]. At all stages of pregnancy, the occurrence of any abnormal situation may lead to serious consequences, jeopardizing the normal development of the fetus and the health of the mother. For example, placental dysfunction [3], maternal nutritional imbalance [4], or endocrine disorders [5] can lead to adverse outcomes, including premature birth, fetal growth retardation, and even miscarriage. Therefore, it is of great importance to monitor the pregnancy process as early, comprehensively, and meticulously as possible.

When the body is stimulated, urine is not controlled by the homeostatic regulation mechanism and can better retain the changes resulting from minor stimuli received by the body [6]. Currently, in clinical monitoring of fetal development, ultrasound examination is the primary method, which can provide information on various aspects of the fetal morphology and structure [7]. However, the monitoring effectiveness of ultrasound examination still has limitations, making it difficult to accurately capture earlier and more detailed embryonic development changes and potential abnormalities.

Urinary proteomics, as an emerging and highly promising research field, has attracted much attention due to its high sensitivity. Numerous studies have confirmed that the urinary proteome can effectively distinguish the occurrence and development of various diseases, including coronary artery disease [8], bladder cancer [9], glioma [10], autism [11], etc. Urine has become an important source of biomarkers for disease diagnosis and monitoring. For example, in the field of pregnancy, existing research has shown that information on embryonic development during pregnancy can be reflected in the urinary proteome [12]. However, the current research on the changes in the urinary proteome during the pregnancy of rats is still not detailed and comprehensive enough.

In this study, female Wistar rats were selected and urine samples were collected from the 1st to the 18th day of pregnancy, and a non - pregnant control group was also included. The aim is to identify the day-by-day differences in the urinary proteome during the pregnancy of rats through comparative analysis, so as to provide more abundant and accurate data support for a deeper understanding of the fetal and maternal physiological mechanisms of pregnancy.

## 2. Materials and methods

### 2.1. Rat caging

The 8-weeks old Wistar rats (8 females and 4 males) were purchased from Beijing Vital River Laboratory Animal Technology Co., Ltd. All rats were housed in a standard environment (room temperature: 22 ± 1 °C, humidity: 65 - 70%). After acclimatization to the new environment for one week, the experiments commenced. Male and female rats were caged at a ratio of 1:1 at 16:00 PM. The next day at 7:00 AM, the female rats were examined for vaginal plugs. The female rats with vaginal plugs were regarded as successfully mated.

All experimental operations complied with the review and approval of the Ethics Committee of the College of Life Sciences, Beijing Normal University, with the approval number CLS-AWEC-B-2022-003.

### 2.2. Urine sample collection

For the 8 female rats, urine was collected for the first time from 20:00 to 8:00 the day before the start of mating, which was recorded as the pre-pregnancy urine sample D0. Then 4 female rats were randomly selected as the pregnancy group, and the remaining 4 female rats as the control group. The female rats in the pregnancy group were caged with the male rats from 16:00 to 7:00 the next day, while the control group was not caged. From 20:00 to 8:00 on the same day, urine was collected for the second time from the 8 female rats. The pregnancy group was recorded as the first-day pregnancy urine sample E1, and the control group was recorded as the first-day non-pregnancy urine sample D1. Thereafter, urine was collected from the pregnancy group and the control group every day from 20:00 to 8:00 and recorded as E2 - E18 and D2 - E18, respectively. All the samples were immediately stored in a −80 °C freezer after collection.

### 2.3. Preparation of urine samples

Rat urine samples were processed with a Magicomics-AP-9 automated platform (Beijing Qinglian Biotech Co., Ltd, China) using the Magicomics DMB assay kit (Beijing Qinglian Biotech Co., Ltd, China) following the manufacturer’s instructions. Briefly, 10 μL of DMB beads were washed with 150 μL of dilution solution, then 50 μL of urine sample and 100 μL of dilution solution were added. The mixture was incubated at 37 ° C for 1 h with shaking at 1000 rpm. After incubation, the supernatant was removed by magnetic separation, and the DMB beads were washed three times with 150 μL of dilution solution. Subsequently, 50 µL of enzyme solution (containing Tris(2-carboxyethyl) phosphine (TCEP) and 2-Chloroacetamide (CAA)) was added to resuspend the DMB beads, followed by digestion with 0.5 µg of trypsin at 37°C for 4h. After incubation, the mixture was placed on a magnetic separator to settle, and 45 µL of supernatant was transferred to a new centrifuge tube. Then, 25 µL of loading solution and 25 µL of stop solution were added and mixed. The entire liquid was loaded onto a desalting column, washed with 100 µL of wash solution 1 and 100 µL of wash solution 2. After changing the centrifuge tube, 100 µL of elution solution was added and the eluate was collected, freeze-dried, and analyzed using LC-MS/MS.

### 2.4. Proteome analysis of urine samples

LC-MS/MS analysis was conducted using a timsTOF HT mass spectrometer (Bruker, Germany) coupled with an UltiMate 3000 liquid chromatography system (Thermo Fisher Scientific, USA). For each sample, 400 ng of peptides dissolved in 0.1% formic acid (FA) aqueous solution were injected. The liquid chromatography conditions included a C18 reverse-phase analytical column (C18, 1.5 μm, 100 μm × 15 cm). Mobile phase A consisted of 0.1% FA, while mobile phase B comprised 80% acetonitrile and 0.1% FA. The gradient was as follows: 0-17 min (3.5-32% B), 17-18 min (32-95% B), 18-20 min (95% B), and 21-22 min (95%-1% B).

All the samples were analyzed in Data-independent acquisition (DIA) mode with a mass scan range of m/z 300-1500, and the primary mass resolution of 60000 (at 1222 m/z). In the TIMS tunnel, an accumulation time of 50 ms was set. The capillary voltage was adjusted to 1.5 kV, and the ion mobility ranged from 0.70 to 1.30 cm2/(V). The total cycle time was 1.23 s. For library construction, 6 fractions of pooled samples were analyzed in Data-dependent acquisition (DDA) mode with a mass scan range of m/z 100-1700, an accumulation time of 100 ms in the TIMS tunnel, the capillary voltage of 1.6 kV, the ion mobility ranging from 0.6 to 1.6 cm2/(V), and the total cycle time of 1.1 s with 10 PASEF cycles.

The raw data were analyzed with Spectronaut software (version 18.1.230626.50606) using the UniProtKB Rat database (2023.03.01 released), and a hybrid library constructed by the fractionated pooled samples. The parameters were set as follows: the protease was specified as trypsin with 2 missed cleavage sites allowed; variable modifications included methionine oxidation (Oxidation (M)) and protein N-terminal acetylation (Acetyl (Protein N-term)); the fixed modification was cysteine carbamidomethylation (Carbamidomethyl (C)); the peptide false discovery rate (FDR) was less than 1%. Data filtering was set to a Q-value of 0.01, and normalization was performed using Local normalization.

### 2.5. Data processing and bioinformatic analysis

After removing values less than 1, the quantitative data of all the samples were normalized using the median value of common proteins which were identified in all samples [13]. Proteins detected in all the rats in the pregnancy group in at least one day were retained for further analysis. The missing values were imputed with the global minimum.

All the data were log2 transformed and analysed by the R package Limma (version 3.58.1) for differential expression analysis between the pregnant group and the normal group across all time points from day 1 to day 18. The thresholds for differential proteins were set as follows: *P* < 0.05, |fold change| ≥ 1.5, and without difference between the two groups at day 0. Enrichment analysis of biological processes for differential proteins at each time point was performed using the hypergeometric distribution. The filtering criteria for enriched terms were: *P* < 0.05, background term number ≥ 3, foreground term number (Count) ≥ 2, and Ratio > 0.1. The enrichment analysis of the entire gestation period (days 1-18) highlighted only the top 10 terms with the most significant *P*-values.

## 3. Results and discussion

In this study, a total of 152 urine samples were collected from rats in the gestation group (n=4) and the control group (n=4) before (day 0) or during the gestation period (days 1-18). After low-abundance protein enrichment by DMB beads, 3,455 proteins were identified in all samples (Table 1, Table S1, Figure 1A), with an average of 2,956 proteins per sample. Compared with other studies using multiple fractionation [14] or extended MS gradient (e.g., 120min) [15], our study achieved the highest number of rat urine protein identifications in only 22 min MS run.

**Figure 1.**
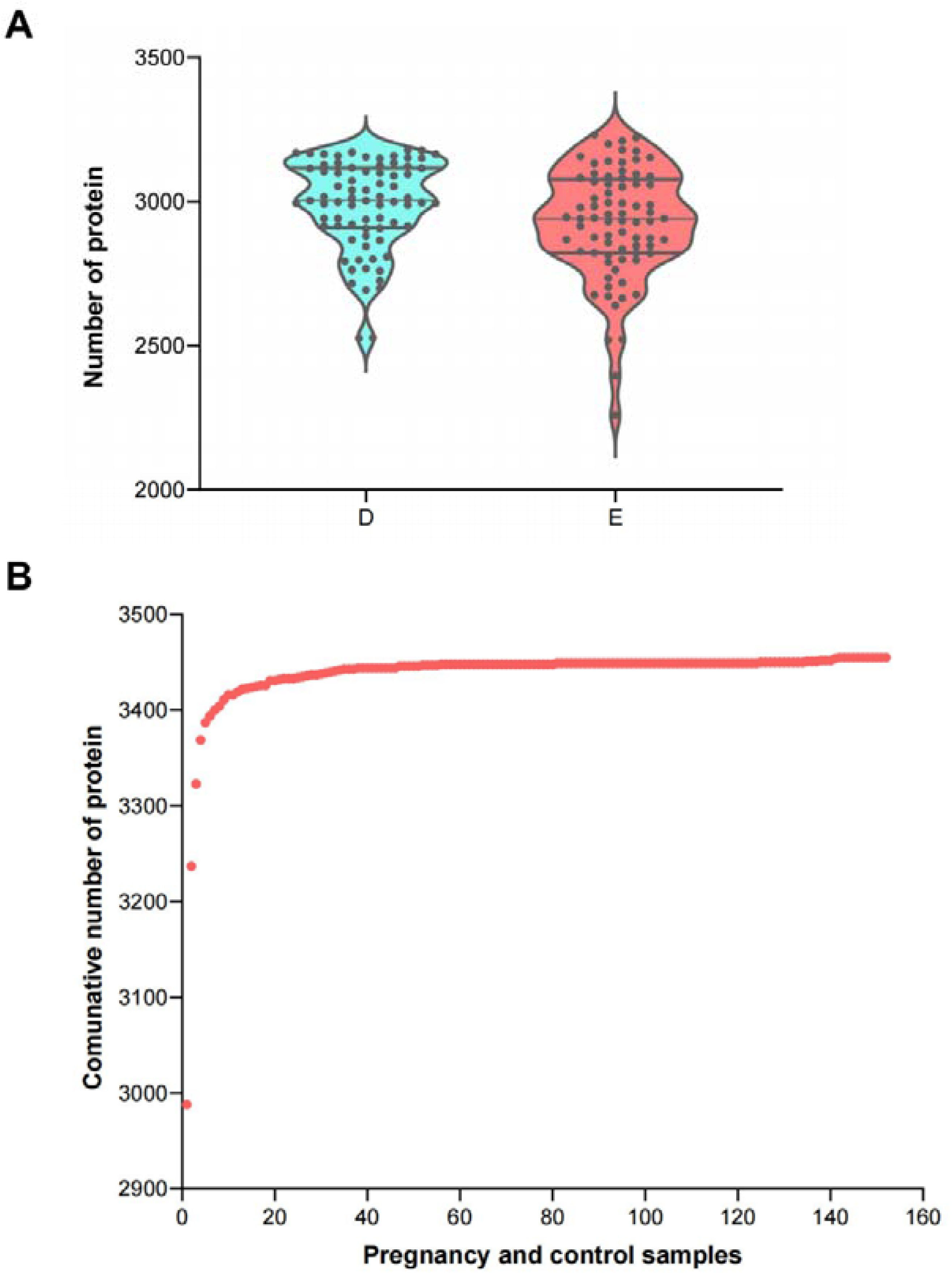
Urine protein identification result. (A) Number of identified urine proteins in each sample (D: control group, E: pregnancy group). (B) Accumulation curves of proteins identified in all the samples.

**Table 1.** Urine protein identification result.

| **Group** | **Number of samples** | **Time point** | **Number of identified proteins** |
| --- | --- | --- | --- |
| Pregnant group | n = 76 | Day 0-Day 18 | 2924 ± 192 |
| Control group | n = 76 | Day 0-Day 18 | 2987 ± 155 |
| All | n = 152 | Day 0-Day 18 | 2956 ± 177 |

### 3.1. The Changes of urine protein in pregnancy rats and control rats

After data filtering, 3,201 proteins were retained in the subsequent analysis. PCA result suggested the difference of urine protein expression between pregnancy group and control group (Figure 2A). Further analysis discovered differentially expressed proteins (DEPs) in urine samples between the two groups from days 1-18 (Figure 2B, Table S2). The number of DEPs showed three peaks: 1-2 days after fertilization, 5-7 days after fertilization (embryo implantation), and 14-18 days after fertilization (before delivery), indicating that there were significant changes in urine proteins during the gestation period, particularly at these three stages. The heatmap also showed the expression changes of all the DEPs from days 1 to 18 after fertilization (Figure 2C).

**Figure 2.**
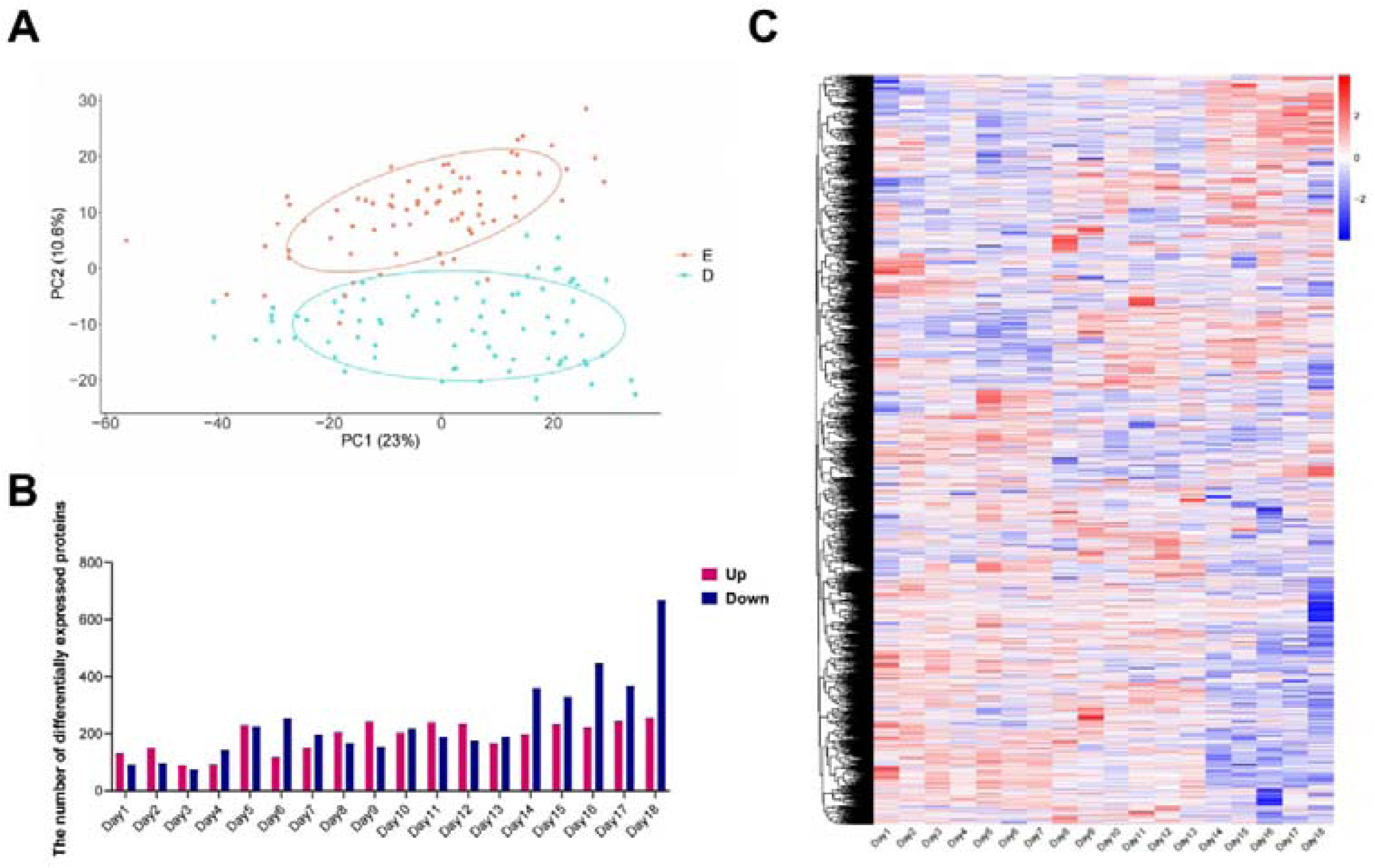
Difference of protein expression between pregnancy group and control group. (A) PCA analysis of all the samples (D: control group, E: pregnancy group). (B) Differentially expressed proteins (DEPs) between pregnancy group and control group at 1-18 days. (C) Heatmap of DEPs between pregnancy group and control group from days 1-18 after fertilization.

### 3.2. The changes occurring throughout the entire gestation period in rats

GO-BP enrichment analysis of DEPs from days 1-18 showed that, compared with control rats, the urinary proteome of pregnant rats gradually exhibited regular up-regulated biological functions during certain stages of pregnancy (Figure 3, Table S3). Corresponding to blastocyst formation and implantation into the uterus after fertilization in early gestation, blastocyst formation was enriched as early as the first day after fertilization. Biological processes associated with cell division were significantly enriched from days 2 to 7, including midbody abscission, mitotic metaphase chromosome alignment, and regulation of centrosome duplication. Additionally, poly-somal sorting-related processes were also enriched during this period, such as positive regulation of exosomal secretion, ubiquitin-independent protein catabolic process via the multivesicular body sorting pathway, and ubiquitin-dependent protein catabolic process via the multivesicular body sorting pathway. Previous studies have demonstrated that MVBs can specifically recognize and mediate the degradation of sperm mitochondria-derived materials (MD) [16].

**Figure 3.**
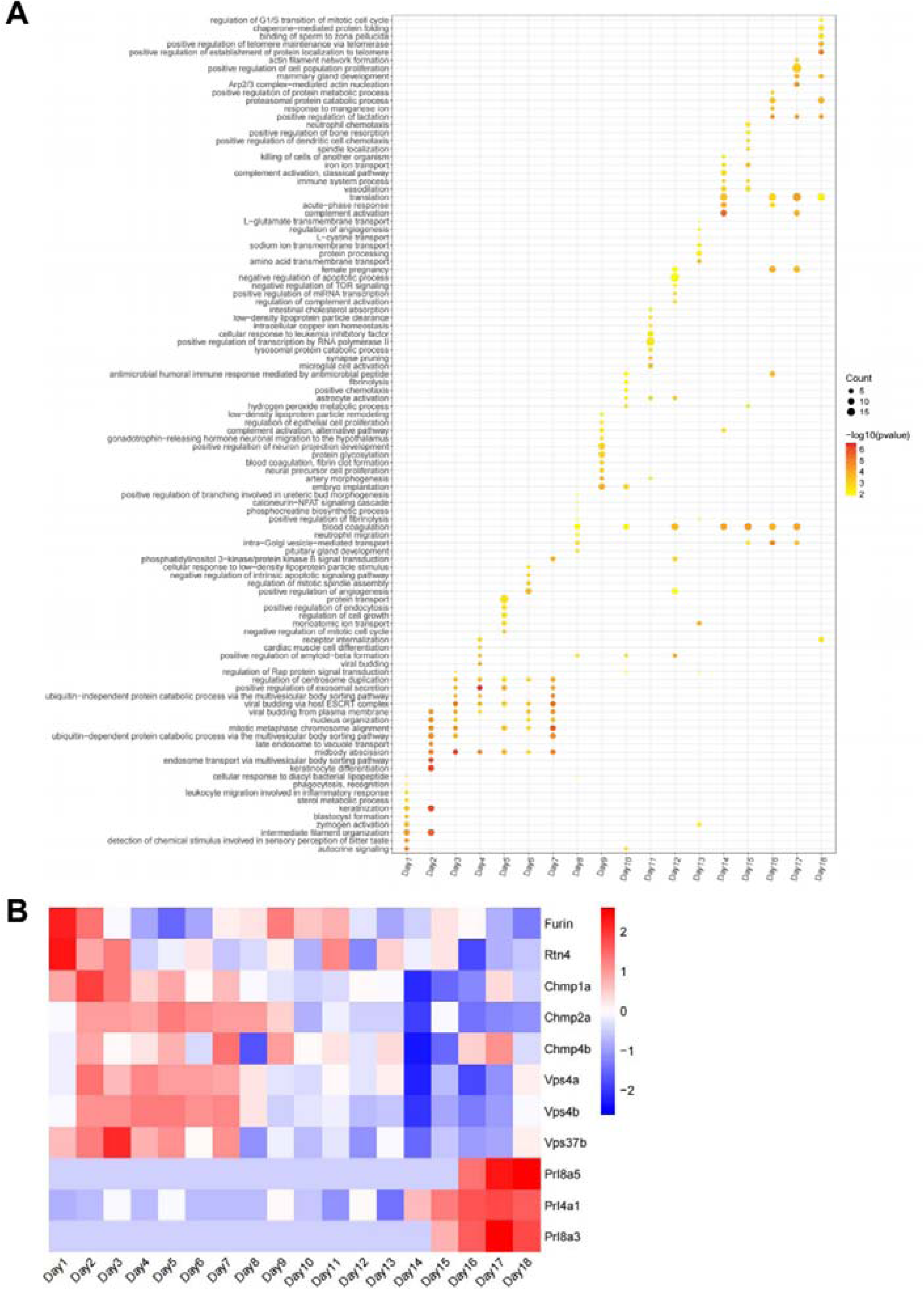
Biological functions (A) and related DEPs (B) of up-regulated rat urinary proteins during the gestation.

In the intermediate and mid-to-late phases of the pregnancy period following embryo implantation, biological processes related to embryo development continuously appeared in the top 10 enriched terms. For example, pituitary gland development and positive regulation of branching involved in ureteric bud morphogenesis were observed on day 8. Artery morphogenesis, neural precursor cell proliferation, positive regulation of neuron projection development, and gonadotrophin-releasing hormone neuronal migration to the hypothalamus were enriched on day 9 of pregnancy. These results may reflect the embryonic development of renal system, vascular system and nervous system during pregnancy [17–19].

Additionally, from the half of gestation, especially in the late of pregnancy, blood coagulation remained active in the pregnancy group. This result is consistent with gradually increased coagulation function from early to late pregnancy [12], which prevent excessive bleeding during delivery. Fibrinolysis increased around day 10, which is consistent with elevated antithrombin time during and after fetal organogenesis to prevent thrombus formation [20]. Concurrently, we observed enrichment of terms related to complement activation during this phase, including complement activation alternative pathway, complement activation-classical pathway, and regulation of complement activation. Complement activation exerts multifaceted functional roles during pregnancy, including protective and destructive actions at the placental level, complement activation at the fetal-maternal interface to defend against pathogens, and aiding in the removal of apoptotic and necrotic cells [21].

At the end of gestation, the stage before delivery, the biological processes associated with lactation were enriched on days 16-18. The mammary gland is a dynamic organ regulated by reproductive and metabolic hormones, developing from puberty and forming a branched, milk-secreting structure at the end of pregnancy. Lactation begins post-placental delivery with progesterone withdrawal, sustained by increased prolactin and oxytocin secretion, and stimulated by infant suckling [22, 23]. This represents a specific change during pregnancy that has not been noted in previous studies of urine proteins in pregnant rats [12].

Changes of key proteins in the biological processes described above were shown in Figure 3B, including proteins Furin and Rtn4 associated with blastocyst formation [24, 25], Chmp family proteins (Chmp1a, Chmp2a, Chmp4b) and Vps family proteins (Vps37b, Vps4a, Vps4b) associated with cell division [26], Prl family proteins (Prl4a1, Prl8a3, Prl8a5) associated with lactation [27].

### 3.3. Dynamic changes of embryonic development throughout the whole pregnancy

#### Embryonic development

Since biological processes related embryo development were continuously enriched in the top 10 terms during pregnancy, we further investigated all enriched terms related to embryonic development throughout days 1-18 of gestation. In addition to blastocyst formation observed on day 1, other biological processes directly linked to embryonic development were also discernible in the rat urinary proteome (Figure 4A, Table S4). For example, events related to gastrula and placenta development, including gastrulation, mesodermal cell differentiation, cell migration involved in gastrulation, and labyrinthine layer blood vessel development. Other events related to morphogenesis of anatomical structure were also observed during pregnancy, including notochord formation, cochlea morphogenesis, embryonic cranial skeleton morphogenesis, embryonic limb morphogenesis, embryonic forelimb morphogenesis, embryonic skeletal system morphogenesis, and embryonic morphogenesis. Post-embryonic development occurred on day 16, whose specific outcome is the completion of embryonic development to the mature structure.

**Figure 4.**
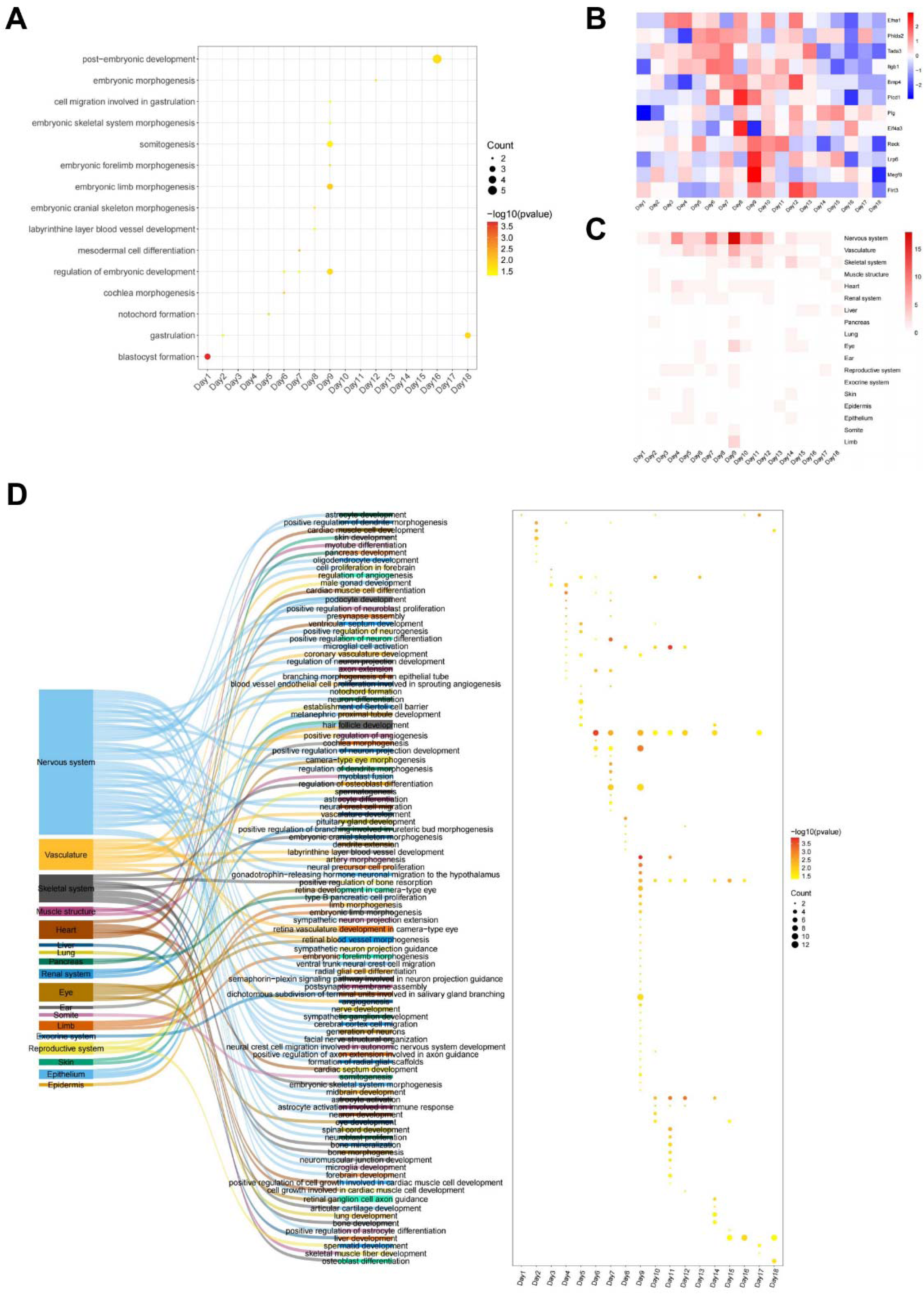
Dynamic changes of enriched biological processes related to embryonic development in pregnant rat urine proteins. (A) Biological processes directly related to embryonic development. (B) Heatmap of proteins related to embryonic development. (C) Biological processes were aligned to the development of 18 organs during the pregnancy period. (D) Specific biological processes related to the development of 18 organs.

Changes of key proteins in the biological processes described above were shown in Figure 4B, including Efna1 associated with notochords formation [28], Phlda2 and Tada3 associated with the regulation of embryonic development [29, 30], Itgb1 and Bmp4 associated with mesodermal cell differentiation [31–33], Plcd1 and Plg associated with the labyrinthine layer blood vessel development [34, 35], Bmp4 and Eif4a3 associated with embryonic cranial skeleton morphogenesis [36, 37], Lrp6, Reck and Megf8 associated with embryonic limb morphogenesis [38–40], Flrt3 and Bmp4 associated with embryonic morphogenesis [41, 42].

#### Organ development of embryo

To explore the developmental status of embryonic organs and tissues throughout gestation, we systematically summarized terms associated with organ and tissue development from days 1 to 18, utilizing keywords pertinent to tissue and organ development, such as organ development, cell differentiation of organ, organ formation, organ morphogenesis. Throughout the entire gestation, we summarized enriched biological processes into the development of 18 distinct embryonic organs and tissues (Figures 4C and 4D), including tissues in nervous system, vascular system, and skeletal system, as well as principal organs such as the heart and liver. Surprisingly, terms related to the nervous system and vascular system were almost present throughout the entire gestation period of rats, particularly those related to the nervous system. Terms related to the development of heart and kidney system were significantly enriched in the early and middle stages of gestation, while those related to liver development were more pronounced in the later stage. Terms related to somitogenesis and limb development were enriched on day 9 of pregnancy. These observations in the urine of pregnant rats indicate that dynamic changes of embryonic development are directly reflected in the alterations of the maternal urinary proteome.

## 4. Conclusion

In this study, daily urinary proteome alterations during rat pregnancy were analyzed. Compared with the control group, changes associated with blastocyst formation and cell division were observed in the early stage of pregnancy. In the intermediate and mid-to-late phases of the pregnancy, changes associated with embryonic development and organ morphogenesis were revealed. Maternal-specific changes, such as lactation, was found in the late stage of pregnancy. These results suggest that both fetal and maternal physiological alterations during pregnancy can be detected in the urinary proteome.

## Supplementary Materials

Supplementary Tables S1-S4.

## Data Availability Statement

The datasets in this study will be available upon acceptance for formal publication (https://www.iprox.org/).

## Author Contributions

Conceptualization, YG, WS and LS. Sample collection: HW. DIA experiments, SC and LS. Proteome data analysis, LG and WS. Analysis results interpretation, LG, WS and YG. Writing—Original Draft, LG, WS, HW and SC, with input from co-authors. Writing—Review & Editing, YG, HW, and LG. All authors listed have made a substantial, direct, and intellectual contribution to the work and approved it for publication.

## Institutional Review Board Statement

The animal study protocol was approved by the Ethics Committee of the College of Life Sciences, Beijing Normal University, with the approval number CLS-AWEC-B-2022-003.

## Informed Consent Statement

Not applicable.

